# Distinct non-coding RNA cargo of extracellular vesicles from M1 and M2 human primary macrophages

**DOI:** 10.1101/2022.08.19.504493

**Authors:** Paschalia Pantazi, Toby Clements, Morten Venø, Vikki M Abrahams, Beth Holder

## Abstract

Macrophages are important antigen presenting cells which can release extracellular vesicles (EVs) carrying functional cargo including non-coding RNAs. Macrophages can be broadly classified into M1 ‘classical’ and M2 ‘alternatively-activated’ macrophages. M1 macrophages have been linked with inflammation-associated pathologies, whereas a switch towards an M2 phenotype indicates resolution of inflammation and tissue regeneration. Here, we provide the first comprehensive analysis of the small RNA cargo of EVs from human M1 and M2 primary macrophages. Using small RNA sequencing, we identified several types of small non-coding RNAs in M1 and M2 macrophage EVs including miRNAs, isomiRs, tRNA fragments, piRNA, snRNA, snoRNA and Y-RNA fragments. Distinct differences were observed between M1 and M2 EVs, with higher relative abundance of miRNAs, and lower abundance of tRNA fragments in M1 compared to M2 EVs. MicroRNA-target enrichment analysis identified several gene targets involved in gene expression and inflammatory signalling pathways. EVs were also enriched in tRNA fragments, primarily originating from the 5’ end or the internal region of the full length tRNAs, many of which were differentially abundant in M1 and M2 EVs. Similarly, several other small non-coding RNAs, namely piRNAs, snRNAs, snoRNAs and Y-RNA fragments, were differentially enriched in M1 and M2 EVs; we discuss their putative roles in macrophage EVs. In conclusion, we show that M1 and M2 macrophages release EVs with distinct RNA cargo, which has the potential to contribute to the unique effect of these cell subsets on their microenvironment.

## Introduction

Macrophages are professional antigen presenting cells present in almost all adult tissues. This heterogenous cell type plays various roles, including defence against pathogens, wound healing, and regulation of other immune cells. Tissue macrophages comprise both resident cells developed during embryogenesis, and recruited macrophages that are sourced through differentiation and extravasation of blood monocytes. Macrophages are often classified into two broad subsets: the M1 ‘classical’ macrophage and the M2 ‘alternatively-activated’ macrophage, which follows the same concept of type 1/2 T cell immunity, and is the most utilised model for the study of macrophage function *in vitro.* M1 macrophages act early against pathogen signals and are involved in inflammatory response, while M2 macrophages have anti-inflammatory properties and help facilitate tissue repair (Reiner, 2009). The M1/M2 paradigm is an oversimplification of the true complexity of macrophage populations *in vivo,* but nonetheless has been a beneficial *in vitro* model in the investigation of macrophage biology, and has been applied to many sites in the human body in which specialised tissue macrophage populations reside, including the liver (Kuppfer cells) (Wan et al., 2014), lungs (alveolar macrophages) (Hu and Christman, 2019) and the placenta (Hofbauer cells) (Reyes and Golos, 2018). The phenotype is not fixed, and M1/M2 cells can switch their phenotype upon changes in their microenvironment (Italiani et al., 2014). The presence of macrophages expressing M1 markers has been linked with inflammation-associated pathologies such as type 2 diabetes, whereas a switch towards an M2 phenotype in tumour associated macrophages is pertinent to more aggressive malignancies (Parisi et al., 2018).

Macrophages both release, and respond to, extracellular vesicles (EVs), the couriers for intercellular communication (Khalife et al., 2019, Singh et al., 2011, Saha et al., 2017, Yang et al., 2021). EV cargo consists of various functional molecules including non-coding RNAs (Veziroglu and Mias, 2020), functional RNA molecules that are not translated into proteins, but regulate gene expression, both *in cis* and *in trans* (Jacob and Monod, 1961). Small non-coding RNAs, which are both present and functional in immune cell EVs, include microRNAs (miRNA), piwi-interacting RNAs (piRNA), Y-RNA, transfer RNA (tRNA), small nuclear RNAs (snRNAs) and small nucleolar RNAs (snoRNAs) (Nolte-’t Hoen et al., 2012). MicroRNAs, and their variants called isomiRs, represent the most thoroughly studied group of small RNAs. They are found in the genome as individual genes or clusters of several miRNAs and control gene expression through several mechanisms including translational repression, mRNA deadenylation and degradation (Hammond, 2015). PiRNAs interact with the PIWI subfamily of Argonaut proteins to preserve genomic stability by repressing the expression of transposable elements, mainly in the germline (Luteijn and Ketting, 2013). Y-RNAs are encoded by four genes in humans and are involved in DNA replication initiation, are structural components of riboprotein complexes, and may also play a role in RNA surveillance and quality control (Kowalski and Krude, 2015). The “house-keeping” tRNAs, snRNAs and snoRNAs, are involved in the RNA maturation and translation of messenger RNA (Shen et al., 2018). While tRNAs carry the building blocks of the newly synthesised protein, tRNA fragments (tRFs), that derive from the cleavage of (pre)tRNAs, were recently discovered and are implicated in several biological/ cellular processes such as gene expression and stress response (Schimmel, 2018). The spliceosome associated snRNAs are involved in splicing and the snoRNAs in the modification of other RNAs (Zhang et al., 2019).

Given their limited number and inaccessibility from healthy tissues, and the requirement for large numbers of cells to generate sufficient EVs *in vitro,* comprehensive unbiased information on primary macrophage EV cargo is lacking. Studies utilising human cell lines, such as THP-1 (Yao et al., 2019), or murine bone marrow-derived macrophages (Bouchareychas et al., 2020) have provided useful insights into macrophage EV cargo, but do not fully recapitulate primary human macrophages due to their immortalisation/requirement for phorbol 12-mytistate-induced differentiation, and species differences, respectively. Most studies involving EVs and macrophages have looked at macrophages as recipients of EVs from various cellular sources. Studies of EV cargo released from human macrophages largely focused on proteomics (Cypryk et al., 2014, Cypryk et al., 2017) or miRNA panels (Roth et al., 2015), while there is also a number of interesting studies that have investigated changes in EV cargo following viral or bacterial infection (Cypryk et al., 2017, Bhatnagar and Schorey, 2007, Singh et al., 2015). Here, we have isolated EVs from M1 and M2 monocyte-derived macrophages from healthy human blood and performed small RNA sequencing to provide a comprehensive insight into the small RNA cargo of human macrophage M1 and M2 EVs, offering an integral dataset for future research.

## Materials and methods

### Monocyte isolation from peripheral blood

Adult human blood was acquired from Research Donors, a HTA licenced and ISO 9001 2015 certified company with Research Ethics Committee (REC) approval (Reference: 20/LO/0325), purchased via Cambridge Bioscience. Signed informed consent was obtained. 240mL peripheral blood was collected from non-fasted healthy female donors (age 21-40 years) into sodium-heparin tubes (Cat. No. 455051, Greiner BioOne) and transported at room-temperature to the laboratory within 6 hours for processing. Blood was centrifuged at 400xG for 10min to deplete platelets, followed by peripheral blood mononuclear cell (PBMC) isolation according to the manufacturer’s instructions for Histopaque-1077 (Cat. No. H8889-500ML, Sigma). The buffy coat was collected, diluted with PBS and centrifuged at 600xG for 10min and 400xG for 10min to further deplete platelets. PBMC were seeded at 1.5×10^6^ cells per cm^2^ in X-VIVO 10 media (Cat. No. BE04-743Q, Lonza) +1% penicillin-streptomycin (Cat. No. P0781, Sigma) on tissue culture plates/flasks, and monocytes allowed to adhere for 1 hour. Non-adhered and weakly attached cells were then removed by vigorous washing with PBS before continuing culture.

### Monocyte-derived M1 and M2 macrophage generation

Macrophage culture was performed under entirely serum-free conditions throughout using X-VIVO 10 media (Cat. No. BE04-743Q, Lonza) supplemented with 1% penicillin-streptomycin (Cat. No. P0781, Sigma). M1 cells were generated by addition of 20ng/mL of granulocyte-macrophage colony-stimulating factor (GM-CSF, Cat. No. 572904, BioLegend) and M2 macrophages with 20ng/mL of macrophage –colony-stimulating factor (M-CSF, Cat. No. 574804, BioLegend). After 6 days, 50% culture volume of fresh media containing GM-CSF or M-CSF was added. On day 7, to complete the polarisation, M1 cultures were treated with 20ng/mL of IFN-γ (Cat. No. 300-02, Peprotech) and LPS from *Salmonella enterica* (Cat. No. L2137, Sigma), and M2 cells were treated with 20ng/mL IL-4 (Cat. No. 200-04, Peprotech) and IL-13 (Cat. No. 200-13, Peprotech) for 48 hours total, with replacement of media and treatments after 24 hours (summarised in Figure 1a).

**Figure 1.**
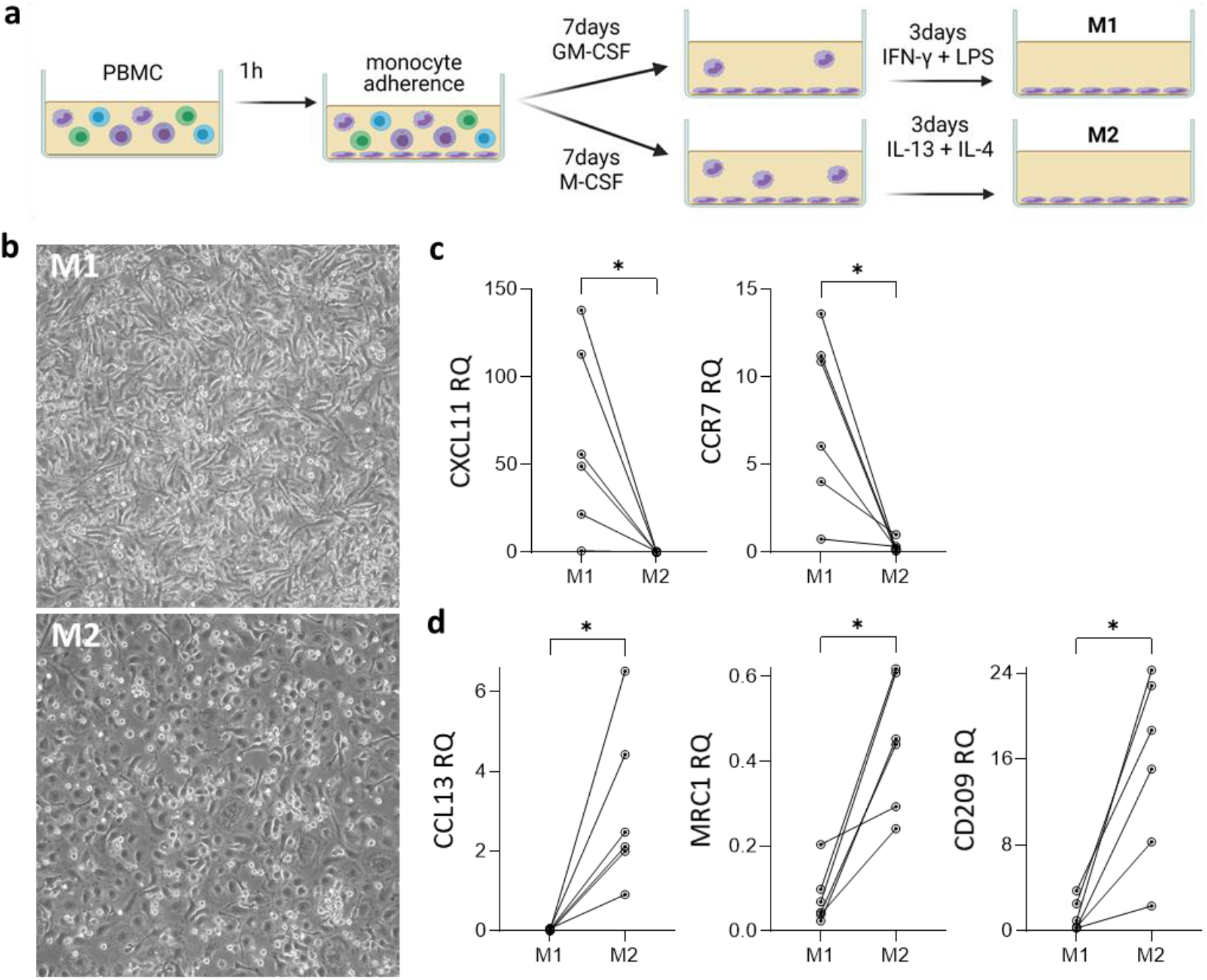
Generation of M1 and M2 monocyte-derived macrophages. a) Monocytes were isolated from PBMC by adherence and differentiated using GM-CSF or M-CSF, followed by IFNγ+LPS or IL-14+IL-13 to create M1 and M2 macrophages, respectively. b) Representative images of M1 and M2 macrophages at 10 days, visualised by light microscopy. c) Gene expression analysis of M1 markers CXCL11 and CCR7 and; d) M2 markers CCL13, MRC1 and CD209 in M1 and M2 macrophages at 10 days measured by qPCR. RQ; relative quantification. * P<0.05; Wilcoxon signed rank test; n=6.

### qPCR for M1 and M2 macrophage phenotyping

Total RNA was isolated from M1 and M2 cells using the Reliaprep^TM^ RNA Cell Miniprep system (Cat. No. Z6010, Promega) following the manufacturer’s instructions, including a DNase I digestion step. RNA was quantified at 260nm using a Nanodrop 2000 spectrophotometer and 300ug were used to create a cDNA library using the High-Capacity RNA-to-cDNA™ Kit (Cat. No. 4387406, Applied Biosystems) following the manufacturer’s instructions. Gene expression was determined using TaqMan Gene Expression Assays containing a FAM dye labelled probe (Applied Biosystems) duplexed with VIC dye labelled probe against GAPDH (Hs99999905_m1, Applied Biosystems) as housekeeping expression control. Probe details are as follows: C-C Motif Chemokine Receptor 7 (CCR7) (Hs01013469_m1), C-X-C Motif Chemokine Ligand 11 (CXCL11) Hs00171138_m1), C-C Motif Chemokine Ligand 13 (CCL13) (Hs00234646_m1), Mannose Receptor C-Type 1 (MRC1) (Hs00267207_m1) and C-Type Lectin Domain Family 4 Member L (CD209) (Hs01588349_m1). The Sso Advanced™ Universal Probes Supermix (Cat. No. 1725281, Bio-Rad) was used to prepare the reactions and real-time qPCR was performed on a STEP-ONE real-time PCR system (Cat. No. 4376357, Applied Biosystems).

### Extracellular vesicle isolation

After polarised M1/M2 macrophage cells were established as above, fresh X-VIVO 10 media with 1% penicillin-streptomycin was added for the 24 hour EV generation period. The conditioned media was collected and centrifuged at 300xG for 5 minutes to remove floating cells, and then at 1000xG for 10 mins to remove remaining cells/debris. The media was concentrated to 500μL using the Vivaspin 15R Hydrasart 30,000 MWCO columns (Cat. No. FIL8452, Sartorius). This 500μL was loaded onto qEV Original (70nm pore) size exclusion chromatography columns (Cat. No. SP1, IZON) in an Automatic Fraction Collector (IZON), and EVs isolated following the manufacturer’s instructions, with PBS used as the buffer. The void volume was 3mL, followed by up to 26x 500μL fractions.

### Nanoparticle tracking analysis (NTA)

EV concentration and size were measured using the ZetaView PMX 120 S (Particle Metrix) with the standard NTA cell assembly installed, operated using the accompanying software ZetaView 8.05.12 SP2 (Particle Metrix). Samples were diluted in 1mL PBS to obtain a concentration within the recommended measurement range (50-200 particles/frame), corresponding to dilutions from 1:100 to 1:250. The instrument captured a 21 second video to measure each sample at 11 different positions, with two readings at each position. After automated analysis of all 11 positions and removal of any outliers, the size and the concentration of the sample were calculated by the software. The instrument pre-acquisition parameters were set to a temperature of 23°C, a sensitivity of 75, a frame rate of 30 frames per second, a shutter speed of 100, and a laser pulse duration equal to that of shutter duration. Post-acquisition parameters were set to a minimum brightness of 30, a maximum size of 1000 pixels, a minimum size of 10 pixels and a trace length of 15. Polymer beads with a uniform size of 100nm (Cat. No. 3100A, Thermo Fisher) were used to calibrate the instrument prior to sample readings.

### Protein assay and silver stain of EV fractions

Protein was isolated from EVs by addition of RIPA buffer (Cat. No. 10010263, Cayman Chemical) supplemented with protease (Cat. No. 04693124001, Roche) and phosphatase (Cat. No. 4906837001, Sigma) inhibitors. Protein was quantified by microBCA assay (Cat. No. 23227, Thermo Fisher), read at 562nm on a Nanodrop spectrophotometer. For silver staining, 9μL of protein from each fraction were mixed with reducing sample buffer (Cat. No. J61337, Alfa Aesar), heated at 95°C for 5min and separated on a 10% Bis-Tris gel (Cat. No. WG1203BOX, Invitrogen). Silver staining was performed using the Pierce^TM^ silver stain kit (Cat. No. 24612, ThermoFisher) following the manufacturer’s instructions.

### Electron microscopy

The EV-enriched fractions (1-3) were combined and concentrated using Amicon Ultra 0.5mL 30kDa cutoff spin columns (Cat. No. 10012584, Fisher Scientific). 300 mesh continuous carbon support copper grids (Cat. No. AGG2300C, Agar Scientific) were glow discharged using a Fischione NanoClean Model 1070 instrument. Six microliters of each concentrated sample were applied directly on the grids for 5 min followed by 2% uranyl acetate for 45 sec. The grids were then air dried and imaged using a Tecnai T12 transmission electron microscope at 120 kVolt.

### Western blotting

EV and cellular protein (20μg for Calnexin and 5μg for GAPDH blots) was mixed with reducing sample buffer (Cat. No. J61337, Alfa Aesar), heated at 95°C for 5min and separated on a Bolt 4-12% Bis-Tris gel (Cat. No. NW04122BOX, Thermo Fisher), alongside a Precision Plus Protein Dual Colour Standard (Cat. No. 1610374, Bio-Rad). Semi-dry transfer of protein to a nitrocellulose membrane (Cat. No. IB23001, Thermo Fisher) was performed with the iBlot 2 gel transfer device (Cat. No. IB21001, Thermo Fisher) using a 7 minute, 20V programme. Membranes were blocked with 1% w/v bovine serum albumin (BSA, Cat. No. A7906, Sigma) and 5% w/v skimmed milk, and probed with 8ng/mL anti-calnexin antibody (Cat. No. 2433S, Cell Signalling) overnight followed by 25ng/mL goat anti-rabbit -HRP (Cat. No. P0448, Agilent) secondary antibody or with 600ng/mL anti-GAPDH directly conjugated with HRP (Cat. No. sc-47724, Santa Cruz). Signal was detected by incubation with ECL Prime western blotting system (Cat. No. GERPN2232, GE Healthcare) and measured on an ImageQuant biomolecular imager (Cat. No. LAS4000, GE Healthcare).

### ELISA

ELISAs were adapted from Webber et al. (2015). High-binding 96-well ELISA plates (Cat. No. 655061, Greiner Bio-One) were coated overnight at 4°C with 25μL of SEC fractions 1 to 5 diluted 1:2 with PBS (Cat. No. LZBE17-512F, Lonza). Wells were washed with 0.05% Tween-20 (Cat. No. P9416, Sigma) in PBS. For intraluminal labelling, the bound EVs were fixed with 4% paraformaldehyde (Cat. No. 20909.290, VWR) for 20 minutes at room temperature, followed by permeabilization with 200μL of 0.1% Triton-X (Cat. No. P9284, Sigma) in PBS for 10 minutes at room temperature prior to staining. Wells were blocked with 1% BSA (w/v) and 0.05% Tween-20 in PBS for 2 hours at room temperature. After washing, 100μL per well of the primary antibodies (CD63 (Cat. No. MCA2142, Bio-Rad) and HLA-ABC (Cat. No. ab70328, Abcam)) or isotype controls were applied at 1mg/mL in 0.1% BSA (w/v) and 0.05% Tween-20 ion for 2 hours at room temperature on a plate shaker set to 400rpm. After washing as above, 100μL per well of anti-mouse HRP-conjugated antibody (Cat. No. P0447, Agilent Dako) was applied at 0.1mg/mL in 0.1% BSA (w/v) and 0.05% Tween-20 in PBS for 1 hour at room temperature on a plate shaker set to 400rpm. After washing as above, TMB substrate (Cat. No. 00-4201-56, Invitrogen) was added for 15 minutes at room temperature and the reaction quenched by addition of Stop Solution (Cat. No. MI20031, Microimmune). Absorbance at 450nm was measured on a Molecular Devices Versamax tuneable plate reader.

### EV RNA extraction

Based on our demonstration that SEC fractions 1-3 were enriched in EVs and low in protein, indicating separation of EVs from soluble protein, we combined these for RNA analysis. Total RNA was isolated from 220μL of the combined SEC fractions, using a Total RNA Purification Kit (Cat No. P4-0058 – 17200, Norgen), following the supplementary protocol for exosomal RNA. Briefly, 660μL RL Buffer were added to the EVs followed by 880μl absolute ethanol. The mixture was then applied on the Mini Spin Columns, washed, DNAse I treated (Cat. No. P4-0098 – 25710, Norgen), and eluted in 30μl elution solution. Low yield samples were subjected to concentration using a SpeedVac (Thermo) at medium speed and no heat. RNA concentration was measured on an Agilent 2100 Bioanalyzer system using an RNA 6000 pico chip (Agilent).

### Library preparation and RNAseq

EV RNA samples were prepared for small RNA sequencing using QIAseq small RNA Library Prep kit (Qiagen) using a minimum of 1.1 ng RNA per sample. The library preparation method counteracts amplification bias by adding unique molecular identifier (UMI) on each molecule before amplification and sequencing, so that uneven PCR amplification can be detected and removed as part of the data analysis. The finished libraries were quality controlled using an Agilent 2100 Bioanalyzer system and quantified by qPCR. Libraries were pooled and sequenced on an Illumina NextSeq500 sequencer by single-end 75bp sequencing.

### Bioinformatic and statistical analysis

The raw data was quality filtered and trimmed by fastx_toolkit, and adaptor sequences were removed using Cutadapt. The reads were collapsed to remove identical UMI containing reads. FastQC was used to ensure high quality sequencing data. Filtered reads were mapped using Bowtie to a list of transcriptomes. First, reads were mapped to tRNA sequences from Genomic tRNA Database (GtRNAdb) allowing one mismatch. Unmapped reads were then mapped to miRNAs from miRBase v22 allowing zero mismatches, but allowing for non-templated 3’ A and T bases. Unmapped reads were then sequentially mapped against other relevant RNA datasets allowing one mismatch: snRNA from RNAcentral, snoRNA from snoDB, Y RNA from refSeq and Gencode, rRNA from refSeq, piRNA from RNAcentral, RNA families in Rfam, mRNA from refSeq, followed by the human genome (hg19). This was done to discover which RNA species were present in the sequencing data. The small RNA expression profiles generated were used for differential expression analysis in R using the DESeq2 package, and volcano plots generated using R.

### Further miRNA and tRNA analysis

MicroRNA cluster analysis was performed using the miRNA cluster definitions by miRbase v.22. Clusters were defined as a set of two or more miRNAs produced from genomic locations within 10 kb in the genome. For inferring the potential regulation of target genes by the miRNA cargo of M1 and M2 EVs, the MIENTURNET (MIcroRNAENrichmentTURnedNETwork) webtool was utilised (http://userver.bio.uniroma1.it/apps/mienturnet/). This webtool uses data from TargetScan or miRtarBase and performs statistical analysis for overrepresentation of miRNA-target interactions; we utilised miRtarBase, which is the most up-to-date tool for validated miRNA-target interactions (Licursi et al., 2019). All significantly different miRNAs were input (adjusted p value<0.05) for M1 and for M2 EV cargo. Following target enrichment, the top ten miRNAs for each EV subset (based on lowest p value) were input for functional enrichment analysis. The tRNA fragments were annotated, using the annotate_tRF function of the R package MINTplates designating tRNA originating sequences with a MINTbase ID, also called tRF label or “License Plates” nomenclature. Furthermore, tRNA sequences are annotated by originating tRNA and fragment type: 5’-half, 5’-tRF, i-tRF, 3’tRF and 3’-half per MINTbase definitions (Pliatsika et al., 2018).

## Results

### Generation of polarised M1 and M2 monocyte-derived macrophages

M1 and M2 macrophages were successfully generated by culture in the presence of GM-CSF followed by IFN-γ and LPS, or M-CSF followed by IL-13 and IL-4, respectively (Figure 1a). They had the expected morphology, with M1 cells being more adherent and dendritic-like, and M2 being more heterogenous, with a proportion of cells displaying a ‘fried egg’ appearance (Bertani et al., 2017)(Figure 1b). Real time qPCR profiling of the cells for previously reported M1 and M2 markers (Martinez et al., 2006) confirmed their polarisation; M1 cells had significantly increased expression of the M1 markers CXCL11 (397x fold) and CCR7 (10.5x fold) compared to M2 cells (Figure 1c), whilst M2 cells had significantly higher expression of the M2 markers CCL13 (406x fold), MRC1 (10.1x fold) and CD209 (8.6x fold) compared to M1 cells (Figure 1d). Based on these data, we proceeded to EV generation.

### Characterisation of M1 and M2 macrophage EVs

M1 and M2 macrophage EVs were characterised, including reference to the Minimal information for studies of EVs 2018 (Thery et al., 2018) and the EV-TRACK database (Consortium et al., 2017). This study is registered on EV-TRACK with reference number EV220120. Nanoparticle tracking analysis (NTA) and protein assay showed that EVs were enriched in the SEC fractions 1-3 following the void, and were separated from the increased protein amounts observed from fraction 8 onwards, indicating the appearance of soluble protein (Figure 2a). This increase in free protein in later fractions was also shown by silver staining (Figure 2b). Based on these data, fractions 1-3 were combined for subsequent analyses, providing a mean EV count of 1.12×10^8^ particles/million starting PBMC for M1 cells and 3.95×10^7^ particles/million starting PBMC for M2 cells (Figure 2c). EVs isolated from M1 cells were slightly but statistically significantly larger than EVs from M2 cells (p=0.007), with an average median (±SD) size of 165.4nm (±11.71) for M1 EVs and 156.4nm (±6.79) for M2 EVs (Figure 2d). A representative size distribution of EVs from primary macrophages is shown in Figure 2e. Transmission electron microscopy confirmed EV phenotype, including an abundance of EVs presenting with the classical ‘cupshaped’ morphology (Lobb et al., 2015) (Figure 2f). In our EV preparations, the canonical EV/exosome markers CD63, HLA-A and GAPDH were enriched in fractions 1-3 as measured by ELISA or western blot (Figure 2g-h), while the absence of the negative EV marker calnexin was assessed by western blotting (Figure 2h).

**Figure 2.**
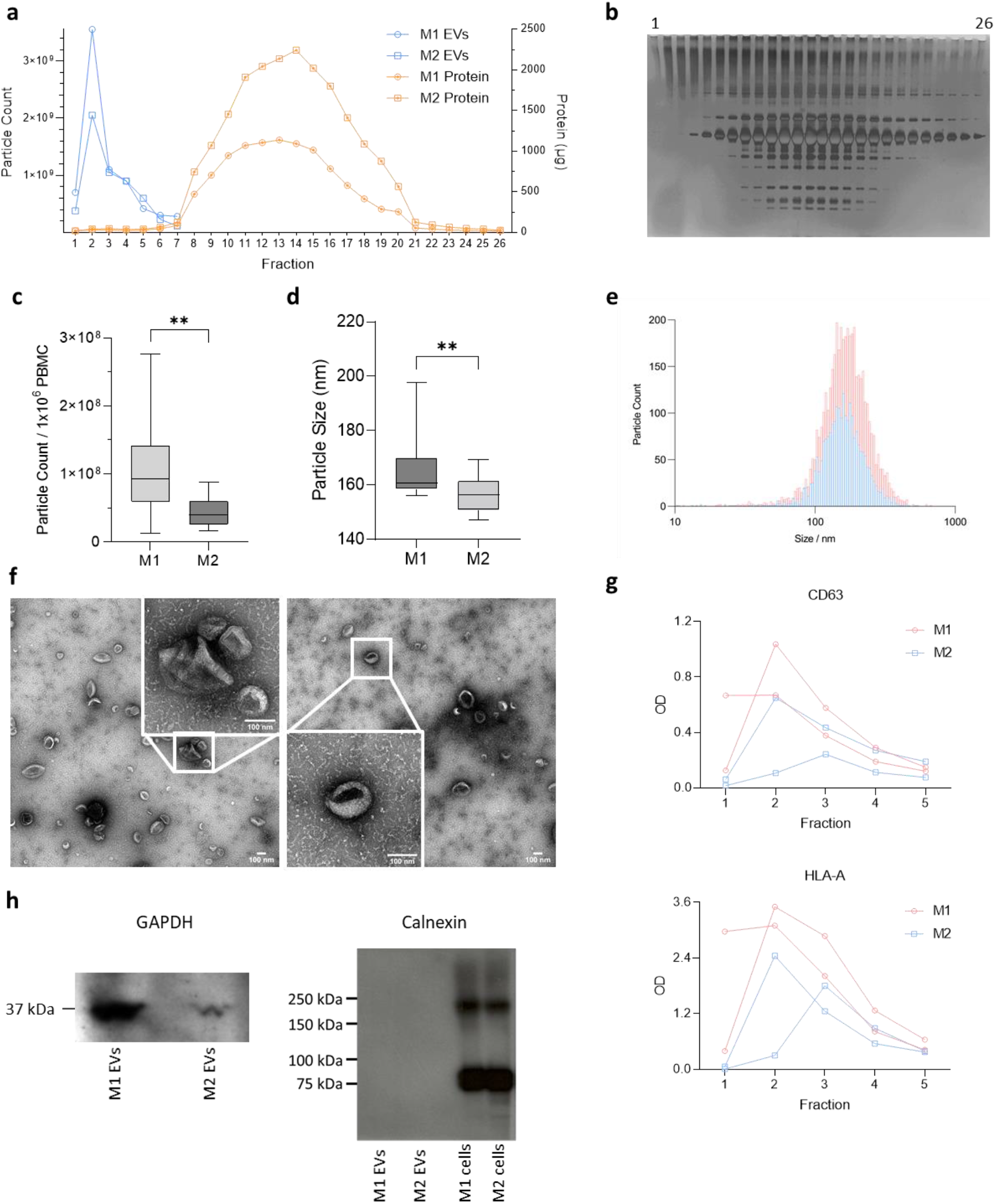
M1/M2 macrophage-derived extracellular vesicle characterisation. Extracellular vesicles were isolated by size exclusion chromatography (SEC) and characterised by nanoparticle tracking analysis (NTA), electron microscopy, western blotting, and ELISA. a) Representative EV count measured by NTA (solid blue line; right Y axis) in the first eight SEC fractions overlaid with the protein concentration in 26 SEC fractions (dashed orange line; left Y axis) showing separation of EVs from soluble protein. b) Representative image of silver staining for the 26 SEC fractions. c) Total particle count measured by NTA in fractions 1-3, per million PBMC seeded. Wilcoxon matched-pairs signed rank test, p≤0.01, n=12. d) Median size of EVs from M1 and M2 cells, measured by NTA. Wilcoxon matched-pairs signed rank test, p≥0.01, n=12. e) Representative size profile of M1 (red) and M2 (blue) macrophage EVs measured by NTA. f) Transmission electron micrographs of EVs from M1 and M2 macrophages. g) ELISA for the canonical EV surface markers CD63 and HLA-A in the first five SEC fractions. h) Western blot for the luminal EV marker GAPDH and the negative EV marker Calnexin.

### M1 macrophage EVs have a higher proportion of miRNAs and lower proportion of tRNA fragments compared to those from M2 macrophages

Sequencing performed on small RNA libraries prepared from EVs produced by M1 and M2 macrophages, detected miRNAs, isomiRNAs, tRNAs, piRNAs, snRNAs, snoRNAs and Y-RNAs (Figure 3a). Classifying the sequenced RNAs revealed several distinct features of the composition of small RNAs and RNA fragments in M1 and M2 EVs (Figure 3a-b). In M1 EVs, miRNAs were the most abundant subclass, making up 58% of the small RNAs, compared to 34% in M2 macrophage EVs (p<0.01; Figure 3b). The next most abundant small RNA subclass was tRNA fragments, which was also significantly differentially abundant in M1 and M2 EVs, making up 45% of the small RNAs in M2 compared to 23% of small RNAs in M1 EVs (p<0.01; Figure 3b). The remaining RNA species, isomiRs, piRNA, snRNA, snoRNA, and Y-RNA made up the 3.2%, 4.3%, 1.3%, 0.2%, and 10% of all small RNAs in M1 macrophage EVs, and the 4.1%, 8.2%, 1.8%, 0.2% and 7.1% of all small RNAs in M2 macrophage EVs, respectively.

**Figure 3.**
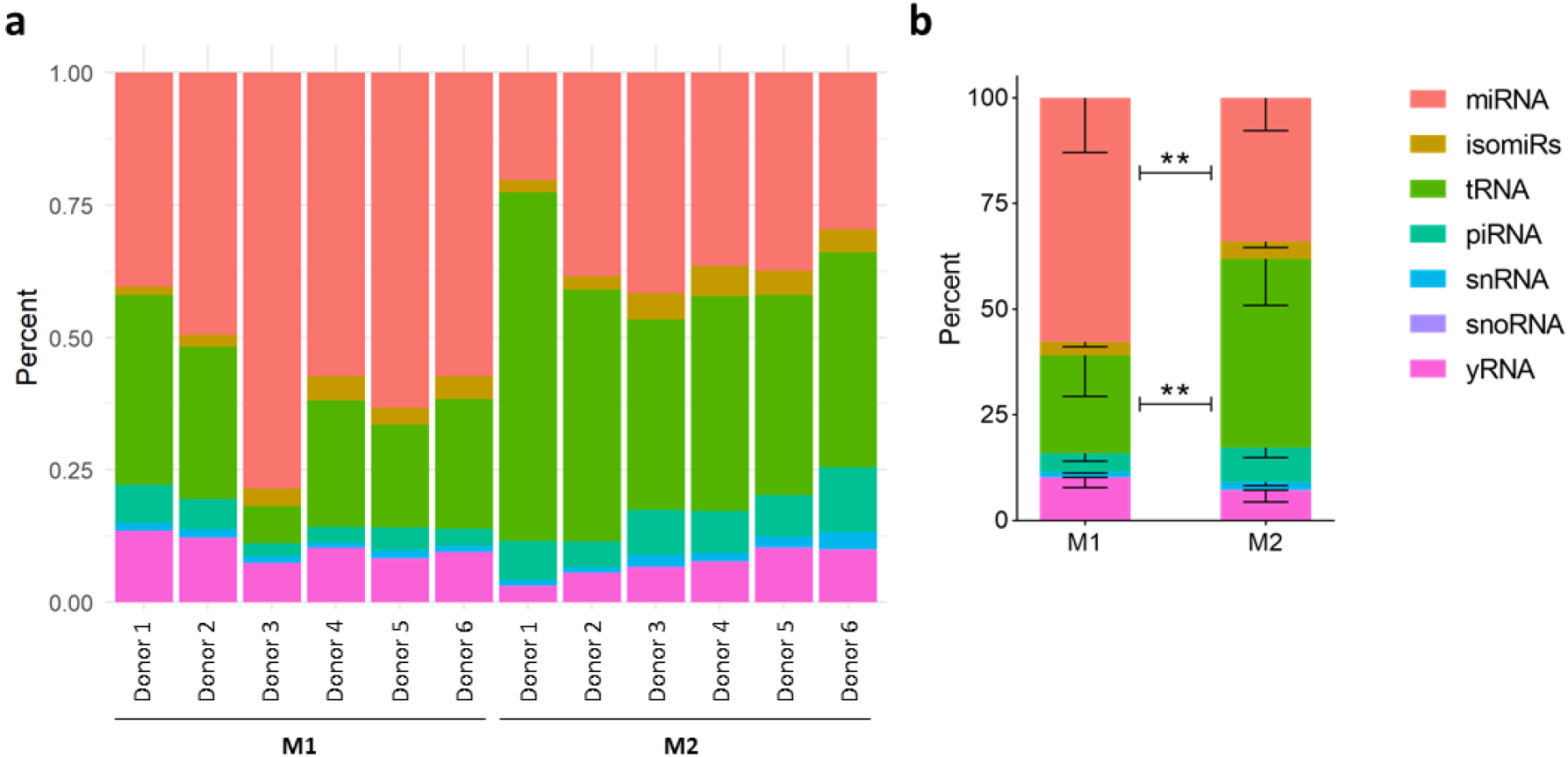
M1 macrophage EVs have higher relative abundance of miRNA and lower relative abundance of tRNA compared to M2 macrophage EVs. a) Relative composition of small RNA species miRNA, isomiRs, tRNA fragments, piRNA, snRNA, snoRNA and yRNA fragments in each sample. b) Comparison between the M1 and M2 EV RNA species. **p<0.01, M1 and M2 compared by repeated measures two-way ANOVA with Geisser-Greenhouse correction, followed by Sidak’s multiple comparisons test.

### Distinct miRNA profiles in EVs from M1 and M2 macrophages

471 miRNAs were detected in M1/M2 macrophage EVs (Supplemental Table 1). The ten most abundant miRNAs in M1 EVs were let-7a-5p, let-7f-5p, let-7i-5p, miR-16-5p, miR-21-5p, miR-26-5p, miR-142-3p, miR-146a-5p, miR-146b-5p and miR-155-5p (Figure 4a). The ten most abundant miRNAs in M2 EVs were let-7a-5p, let-7b-5p, let-7i-5p, miR-16-5p, miR-21-5p, miR-92a-3p, miR-142-3p miR-146a-5p, and miR-486-5p (Figure 4a). Seven of these top ten were shared between the EVs from the two cell types (let-7a-5p, let-7f-5p, let-7i-5p, miR-16-5p, miR-21-5p, miR-142-3p, miR-146a-5p). In all cases, these were more abundant in M1 EVs compared to those from M2, reaching statistical significance for let-71-5p and miR-146a-5p (Supplemental Table 2). Of the 471 miRNAs found in macrophage EVs, 72 were significantly differentially abundant between M1 and M2 macrophage EVs (Supplemental Table 2; Figure 4b), with 43 higher in M1 EVs, and 29 higher in M2 EVs (Figure 4b). MicroRNAs from M1 and M2 macrophage EVs form separate clusters by principal component analysis (Figure 4c).

**Figure 4.**
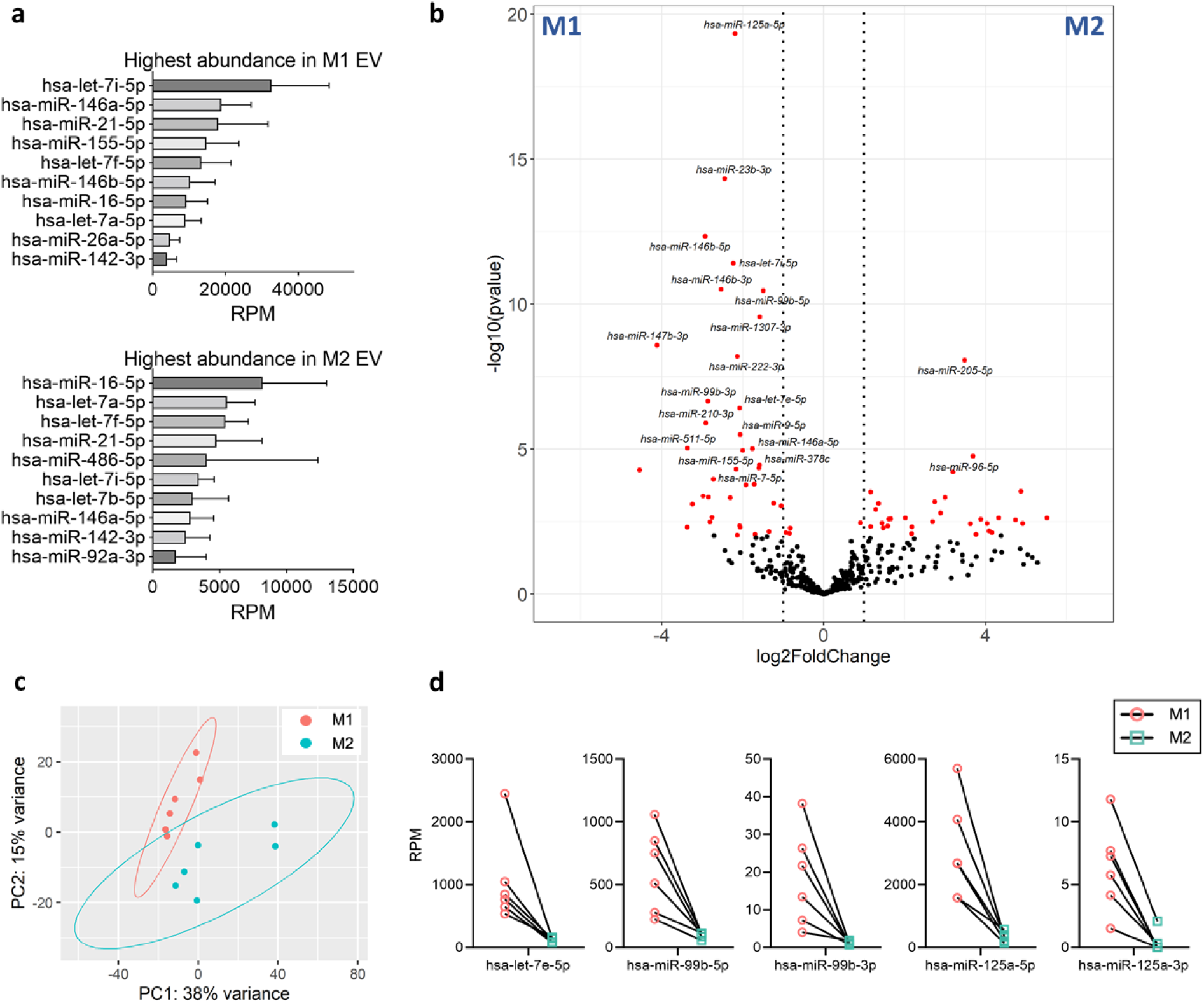
Distinct miRNA profiles in EVs from M1 and M2 macrophages. a) The top 10 most abundant miRNAs in M1 and M2 EVs. RPM; reads per million mapped reads. b) Volcano plot comparing miRNAs between M1 and M2 EVs. X axis shows the log2 fold change and the y axis the log10 p value. Significantly different miRNA (false discovery rate (FDR)<0.05) are coloured red. The twenty miRNAs with the lowest p value are labelled. c) Principal component analysis of miRNAs, with M1 EVs colour-coded red/pink and M2 EV samples colour-coded turquoise. d) The relative abundance expressed as reads per million reads (RPM) of the members of the mir-99b/let-7e/mir-125a cluster in M1 and M2 EVs.

We report all members of the let-7 family in macrophage EVs, except miR-202; with let-7a, let-7f and let-7i being within the top ten most abundant miRNAs in both M1 and M2 EVs. Let-7d-5p, let-7e-5p, let-7i-3p and let-7i-5p were significantly higher in M1 EVs, whilst let-7d-3p was significantly higher in M2 EVs. Finally, miRNA profiles were analysed for miRNA clusters, defined as two or more miRNAs produced from genomic locations within 10 kb. Of the 159 clusters reported by miRbase (v22), 35 clusters were found in M1/M2 macrophage EVs (data not shown). Of these, only one cluster - the miR-99b/let-7e/miR-125a cluster - was notably different between M1 and M2 EVs, with all three miRNAs present in M1/M2 EVs and all miRNAs being significantly more abundant in M1 EVs compared to M2 (Figure 4d). Moreover, both the 5’ and 3’ forms of miR-125a and miR-99b were present and differentially abundant.

### Functional enrichment analysis reveals contrasting targets of M1 and M2 miRNA EV cargo

The gene targets of the miRNA cargo within M1 and M2 EVs were identified using the MIENTURNET webtool (Licursi et al., 2019), looking for overrepresentation of miRNA-target interactions, based on validated miRNA-target interactions from miRTarBase. The top 20 targets of M1 and M2 EVs are shown in Figure 5a-b, and the individual miRNAs targeting these genes are listed in Supplemental Table 3. The top 20 M1 EV miRNA targets include cytokines/chemokines (Interleukins 6 and 8 (IL-6/8) and genes involved in protein trafficking (Sortilin 1 (SORT1), RP2 Activator of ARL3 GTPase (RP2)) and cell proliferation (Microtubule Associated Scaffold Protein 1 (MTUS1), Maternally Expressed 3 (MEG3), A-Kinase Anchoring Protein 8 (AKAP8)). Of note, the Mechanistic Target Of Rapamycin Kinase (MTOR) is one of the top targets for M1 EV miRNA cargo, being targeted by five miRNAs. The top 20 M2 EV miRNA targets include genes involved in transcriptional regulation (Mitogen-Activated Protein Kinase (MAP3K3), Mortality Factor 4 Like 1 (MORF4L1), B-TFIID TATA-Box Binding Protein Associated Factor 1 (BTAF1), SIN3 Transcription Regulator Family Member B (SIN3B), MYC Proto-Oncogene (MYC), Forkhead Box O3 (FOXO3)), oxidative stress (Nth Like DNA Glycosylase 1 (NTHL1), Paraoxonase 2 (PON2)), and NFκB signalling (MAP3K3 and TNF Receptor Associated Factor 4 (TRAF4)). MYC is a top gene target for both M1 and M2 miRNA EV cargo, with ten of the miRNAs upregulated in M1 EVs targeting this gene, compared to eight of those in M2 EVs (Supplemental Table 3). Furthermore, top targets of both also include genes involved in regulation of MYC activity; M1 EV miRNAs target Far Upstream Element Binding Protein 1 (FUBP), which can activate and repress MYC transcription, whilst M2 EV miRNAs target SIN3 Transcription Regulator Family Member B (SIN3B) which represses MYC-responsive genes, and FOXO3 which can prevent MYC translation.

**Figure 5.**
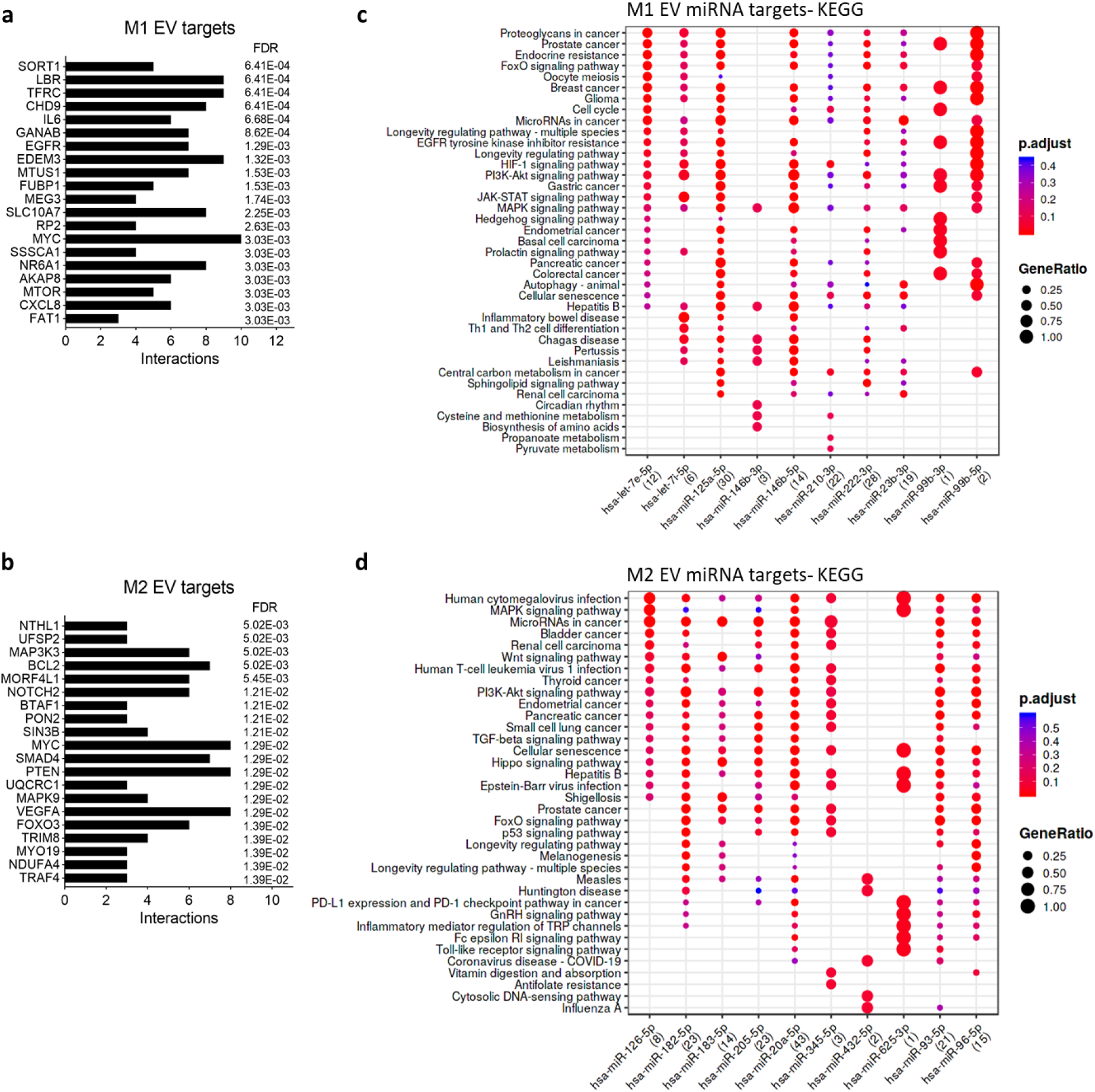
Functional enrichment analysis of targets of miRNAs upregulated in EVs from M1 and M2 macrophages. The miRNAs significantly different between M1 EVs (n=43) and M2 EVs (n=29) were input for MIENTURNET enrichment analysis, based on miRTarBase validated target prediction. a,b) The top 20 gene targets of the miRNA panel significantly upregulated in M1 EVs (a) and M2 EVs (b). Targets ordered by false discovery rate (FDR), bars indicate the number of miRNA interactions for each target. c,d) The top 10 miRNAs that were most significantly differentially abundant, and still present following target enrichment, were then submitted for KEGG pathway analysis. The X axis shows these ten miRNAs, with the number of gene targets in parentheses. The colours of the dots represent the adjusted p-values (FDR), and the size of the dots represents the gene ratio (the number of miRNA targets found annotated in each category divided by the total number of recognised gene targets).

KEGG functional enrichment analysis using the top ten miRNAs present following the target enrichment above identified several interesting biological pathways (Figure 5c-d). For both M1 and M2 EV miRNA cargo, this included potential regulation of the forkhead box (FoxO), the phosphoinositide-3-kinase–protein kinase B/Akt (PI3K/Akt), and mitogen-activated protein kinases (MAPK) signalling pathways. M1 EV miRNA cargo, additionally potentially regulates the JAK-STAT and HIF-1 pathways, as well as autophagy and the cell cycle (Figure 5c). M2 EV miRNA cargo additionally potentially regulates TGF-β, Hippo, p53, GnRH, and Toll-like receptor signalling pathways, as well as senescence, and cytosolic DNA sensing (Figure 5d).

### Distinct profiles of tRNA fragments in EVs from M1 and M2 macrophages, including higher proportion of 5’-halves and 5’-tRFs in M1 EVs

Transfer RNA fragments were classified using MINTmap annotations (Loher et al., 2017), identifying 22,600 tRFs in M1/M2 macrophage EVs (Supplemental Table 4). We classified tRFs into five structural categories by their derivation from full-length mature tRNAs: 5’ halves (5’-half), 3’ halves (3’-half), shorter sequences from the 5’ region (5’-tRF), shorter sequences from the 3’ region (3’-tRF), and those from the internal region (i-tRF) (Kumar et al., 2016)(Figure 6a). The vast majority of tRNAs present in both M1 and M2 EVs were i-tRFs (90.3%; 20,479), followed by 5’tRFs (5.8%; 1,307) and 5’halves (1.5%; 261). Only two 3’-tRFs were identified (the tRF-16-K69YRVD in M2 EVs and the tRF-16-KQ7871B in M1 EVs – nomenclature based on MINTbase ID), and no 3’-halves. M1 macrophage EVs contained a higher number of 5’-halves (1.3x fold, p=0.046) and 5’-tRFs (1.4x fold, p=0.024) compared to M2 macrophage EVs (Figure 6b). 34 tRFs were significantly more abundant in M1 macrophage EVs, and 32 were more abundant in M2 macrophage EVs (Figure 6c; Supplemental Table 5). Of the differentially abundant tRFs, most were again i-tRFs (70% in M1 and 63% in M2), but there was a higher representation of 5’tRFs (18%/39%) and 5’-halves (12%/13%) compared to all identified tRFs. M1 EVs were more enriched in tRFs deriving primarily from tRNA-Leu (n=6), tRNA-Asp (n=6), tRNA-His (n=6), and tRNA-Val (n=4), while M2 EVs were more enriched in fragments from tRNA-Glu (n=13), tRNA-Asp (n=6), and tRNA-Arg (n=5).

**Figure 6.**
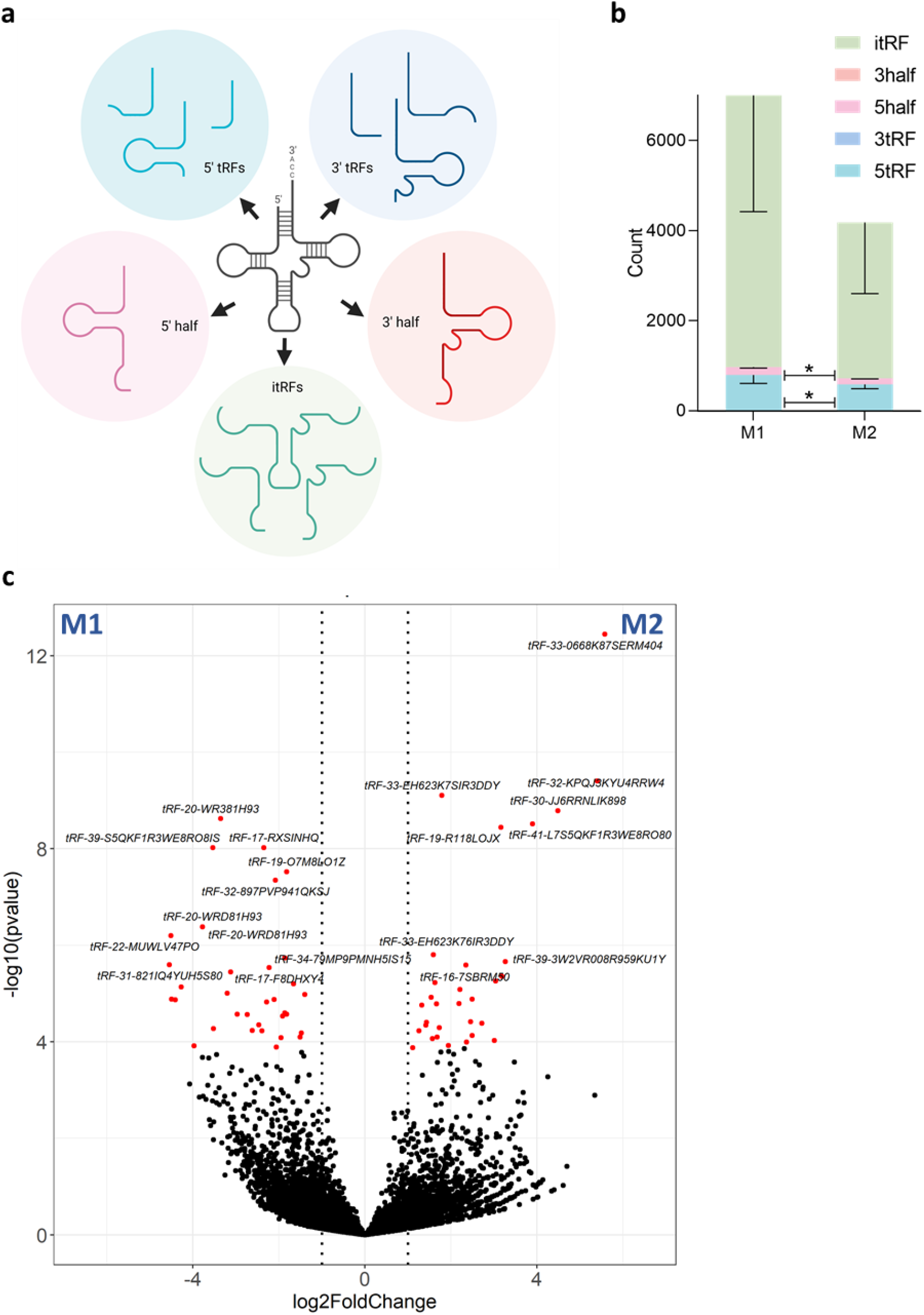
tRNA profiles in EVs from M1 and M2 macrophages. a) tRNA fragments were classified into five types following MINTbase terminology; tRNA halves from the 3’ or 5’ region (3’half/5’half respectively), shorter sequences from the 3’ or 5’ region (3’tRF/5’tRF respectively), or tRNA fragments from the internal region (itRFs). b) The number of all tRNA fragments of each subtype present in EVs from M1 and M2 macrophages. *p<0.05, repeated measures two-way ANOVA with Geisser-Greenhouse correction, followed by Sidak’s multiple comparisons test. c) Volcano plot comparing tRNA fragments between M1 and M2 EVs. The X axis shows the log2 fold change and the y axis the log10 p value. Significantly different tRNA (false discovery rate (FDR)<0.05) are coloured red. The twenty tRNA fragments with the lowest p value are labelled.

### piRNA, snRNA, snoRNA and Y-RNA cargo of M1 and M2 macrophage EVs

A total of 7,059 piRNAs were identified in M1/M2 macrophage EVs (Supplemental Table 6), with 147 of these detected in all M1 EV samples and 74 in all M2 samples. The top two most abundant piRNAs were the same in M1 and M2 EVs-piR-33151 and piR-44757-with a further four found in the top ten of both EV subsets (piR-32017, piR-33151, piR-41603, piR-44757, piR-49143, piR-49144). Six piRNAs were differentially abundant between M1 and M2 EVs (piR-49645, piR-36041, piR-53542, piR-57947, piR-36038, piR-31355); all of which were increased in M1 EVs compared to M2 EVs (Figure 7a; Supplemental Table 6).

**Figure 7.**
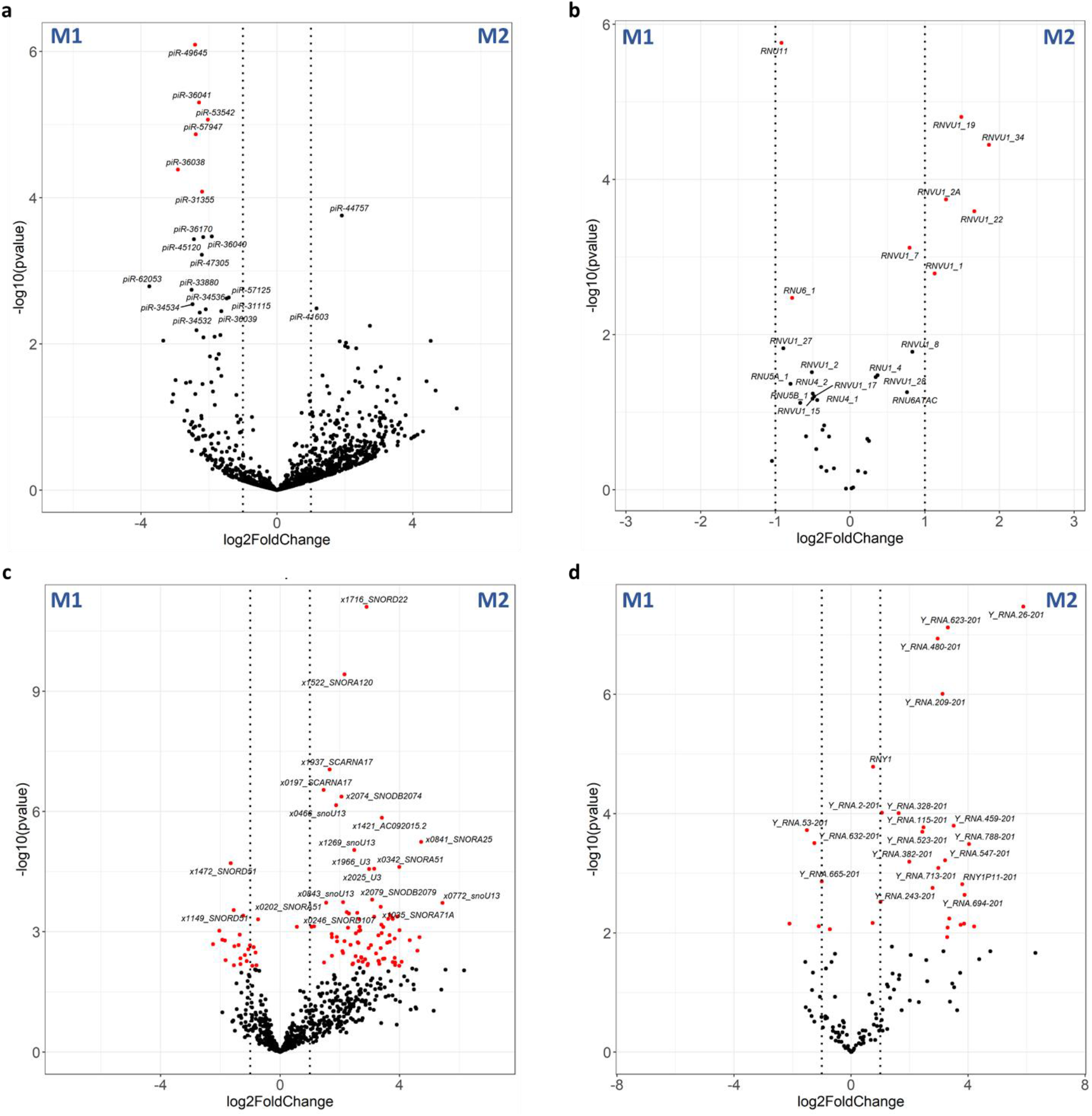
piRNA, snRNA, snoRNA, and yRNA profiles in EVs from M1 and M2 macrophages. a) Volcano plot comparing piRNA fragments between M1 and M2 EVs. b) Volcano plot comparing snRNA fragments between M1 and M2 EVs. c) Volcano plot comparing snoRNA fragments between M1 and M2 EVs. d) Volcano plot comparing yRNA fragments between M1 and M2 EVs. For all volcano plots, the X axis shows the log2 fold change and the y axis the log10 p value. Significantly different tRNA (false discovery rate (FDR)<0.05) are coloured red. The twenty RNAs with the lowest p value are labelled.

45 snRNAs were identified in M1/M2 macrophage EVs (Supplemental Table 7). The most abundant snRNA was the same in both M1 and M2 EVs - RNU2-1 - with a further five found in the top ten of both EV subsets (RNU1-4, RNVU1-28, RNVU1-31, RNU4-1, RNU5B-1, RNVU1-7). Eight snRNAs were differentially abundant between M1 and M2 EVs (Figure 7b; Supplemental Table 7). Two of these (RNU11 and RNU6-1) were more abundant in M1 EVs, and six (RNVU1-19, RNVU1-34, RNVU1-2A, RNVU1-22, RNVU1-7, RNVU1-1) were more abundant in M2 EVs.

1,491 snoRNAs were identified in M1/M2 macrophage EVs (Supplemental Table 8), with 434 of these detected in all M1 EV samples and 114 in all M2 samples. 107 snoRNAs were differentially abundant between M1 and M2 EVs, with 23 more abundant in M1 and 84 more abundant in M2 EVs (Figure7c). SnoRNAs are divided into three subclasses; C/D box snoRNAs (SNORD), H/ACA box snoRNAs (SNORA), and small Cajal body-specific RNAs (SCARNA) (Xie et al., 2007). The most differentially abundant snoRNAs were SNORDs (n=42), followed by SNORAs (n=34) and then SCARNAs (n=7). Of the differentially abundant SNORDs, half were more abundant in M1 EVs, and half were more abundant in M2 EVs (n=21 for each), whereas all but one of the SNORAs (SNORA73B) were higher in M2 EVs, and all the SCARNAs were higher in M2 EVs.

631 Y-RNA derived small RNAs were identified in M1/M2 macrophage EVs (Supplemental Table 9), with 106 of these detected in all M1 EV samples and 28 in all M2 samples. 31 Y-RNAs were differentially abundant between M1 and M2 EVs, with 6 higher in M1 and 25 higher in M2 EVs (Figure 7d).

## Discussion

Here, utilising fully polarised human monocyte-derived macrophages, cultured under serum-free conditions, we report the comprehensive small RNA cargo of EVs from M1 pro-inflammatory macrophages and M2 pro-resolution macrophages; the most common model of human tissue macrophages. Broadening existing knowledge on macrophage RNA cargo, which has hereto focused largely on miRNAs, we demonstrate that M1 and M2 cells produce EVs with both common and disparate miRNA, tRNA, piRNA, snRNA, snoRNA and Y-RNA cargo. Our identification of multiple EV small RNAs specifically associated with pro-inflammatory and pro-resolution macrophage phenotype furthers our mechanistic understanding of macrophage modulation of surrounding cells and tissues.

We found that M1 macrophages release EVs of a larger size than M2 macrophages. The biological significance of this is unclear; potentially reflecting an increased production of larger microvesicles from these cells compared to smaller exosomes. We are currently investigating the proteomic cargo of M1/M2 EVs, which could help answer this question. The apparent increased amount of EVs released from M1 cells could be attributed to the higher cell numbers consistently seen by the end of the polarisation protocol in M1 cultures, likely due to greater adhesion of these cells to the culture plastic (as shown in Figure 1b). It was only possible for us to report EV concentrations relative to seeding cell numbers, as following polarisation, we found that it was not possible to successfully detach the cells without excessive loss, which precluded calculating final cell numbers.

Looking at the overall relative abundance of small RNA cargo in all donors, the most abundant type was miRNA, followed by tRNA fragments, as previously reported for EVs (O’Brien et al., 2020). We observed a higher proportion of miRNA and smaller proportion of tRFs in M1 EVs compared to M2 EVs. Macrophage differentiation in the presence of TLR-4 ligand lipopolysaccharide (LPS), as used in our M1 protocol, can promote miRNA biosynthesis through upregulation of full-length Dicer (Curtale et al., 2019), although notably, this same study also reported equal upregulation of Dicer in human monocytes differentiated to M1 and M2. We detected a significant relative decrease in tRFs in M1 EVs, which could simply reflect the relative increase in miRNAs in these EVs, as tRNA was the second most abundant RNA type, and we saw equal numbers of tRNA fragments increased in M1 and M2 EVs.

MicroRNAs are important regulators of macrophage activation and polarisation (Curtale et al., 2019). We identify a range of miRNA cargo (>400 unique miRNAs) within EVs released from M1 and M2 macrophages, with distinct clustering of M1 and M2 EV miRNA profiles. Of the 72 differentially expressed miRNAs, several have been associated with macrophage polarisation (Zhang et al., 2013), including upregulation of miR-155 and miR-181a in M1 EVs. However, we also found that EV profiles did not always reflect previously reported cellular changes; for example we found higher miR-143-3p and miR-145-5p in M2 EVs, in agreement with reported cellular changes, but significantly lower miR-125-5p and miR-146a-3p, in disagreement (Zhang et al., 2013). This could reflect differences between human monocyte-derived macrophages (mo-macrophages) studied here, and murine bone marrow-derived macrophages studied by Zhang et al., or could indicate selective shuttling of miRNAs from cells into EVs.

The miR-155 is one of the most studied miRNAs in macrophages, and within the immune system in general; widely reported to increase under inflammatory conditions, and decrease under proresolution conditions (Alivernini et al., 2017). Although we indeed report higher expression of miR-155-5p in M1 EVs, several other miRNAs had a greater difference between M1 and M2 EVs. The miRNA with the highest increase (>20x fold-change) in M1 EVs - miR-187-3p - was previously reported to be upregulated in monocytes and mo-macrophages following LPS exposure in an IL-10-dependent manner (Rossato et al., 2012).

The highly conserved let-7 family is among the most abundantly expressed miRNAs in cells, and we report all members of this family in macrophage EVs (except miR-202), with let-7a/let-7f /let-7i being in the top ten most abundant miRNAs in both M1 and M2 EVs. The let-7 family play roles as tumour suppressors, in metabolic reprogramming and in immune system development (Roush and Slack, 2008). In macrophages, let-7c (Banerjee et al., 2013) and let-7b (Wang et al., 2016) are involved in macrophage polarisation, though we find these to be similarly abundant in M1 and M2 EVs. LPS treatment of human monocyte/macrophages upregulates the let-7 containing miRNA cluster miR-99b-5p/let-7e-5p/miR-125a-5p (Basavarajappa et al., 2020, Curtale et al., 2018). We report here for the first time that this cluster is also released by macrophages via EVs; indeed, it was the sole miRNA cluster significantly increased in EVs from M1 macrophages. Release of this cluster via EVs could be a mechanism by which M1 macrophages induce M1 polarisation in bystander cells, in addition to known soluble factors such as cytokines.

Given the large number of miRNAs significantly different between M1 and M2 EVs, we performed target enrichment analysis (Licursi et al., 2019) to investigate the targets of the complete EV miRNA cargo. Top targets of M1 EV-miRNA cargo included IL-6 and IL-8. and several genes involved in cell proliferation, such as the Microtubule Associated Scaffold Protein 1 (MTUS1), the Maternally expressed 3 (MEG3) and the A-Kinase Anchoring Protein 8 (AKAP8). MEG3 is a long non-coding RNA (lncRNA), which modulates the TGF-β pathway (Mondal et al., 2015), and TGF-β regulates macrophage activation, cytokine production and chemotaxis (Murray et al., 2014). Interestingly, Mechanistic Target Of Rapamycin Kinase (MTOR) is targeted by five of the top 20 M1 EV miRNAs. This serine/threonine protein kinase is a central regulator of cellular metabolism, growth and survival in response to hormones, growth factors, nutrients, energy and stress signals (Maiese, 2020). Together with numerous other interesting targets, miRNAs found in both M1 and M2 EVs target the MYC Proto-Oncogene (MYC), required for M2 polarisation (Pello et al., 2012). Thus, the EV miRNA cargo reflects the phenotype of the source macrophage, and represents potential mediators of functional effects in surrounding cells and tissues.

KEGG functional enrichment analysis identified that M1 and M2 EV cargo could potentially regulate the FoxO, PI3K/Akt, and MAPK signalling pathways. FoxO regulates many cellular processes such as cell cycle, apoptosis, metabolism and oxidative stress and immune regulation, and is regulated by the PI3K/Akt pathway (Matsuzaki et al., 2003). The PI3K/Akt pathway also mediates numerous cellular functions including angiogenesis, metabolism, growth, proliferation, survival, protein synthesis, transcription, and apoptosis (Hemmings and Restuccia, 2015). Finally, the equally complex MAPK signalling pathway regulates cell cycle and proliferation, and plays a key role in the immune system (Dong et al., 2002). In addition to these pan-macrophage miRNA-EV targets, M1 EV miRNA cargo additionally regulates the JAK-STAT and HIF-1 pathways, involved in immunity and hypoxic responses respectively. M2 EV miRNA cargo additionally potentially regulates TGF-β and Toll-like receptor signalling pathways, and cytosolic DNA sensing; all important for macrophage responses to pathogen encounter.

In agreement with previous studies on dendritic cells (Nolte-’t Hoen et al., 2012) and T cells (Chiou et al., 2018), we report here the relative high abundance of tRFs in macrophage EVs. As shown in EVs from other cell types (Chiou et al., 2018, Cooke et al., 2019), M1 and M2 macrophage EVs contained primarily fragments of the 5’ end i.e., 5’halves and 5’ tRFs, as well as itRFs. Natural or synthetic 5’ tRFs and not 3’-tRFs can initiate a stress response involving the assembly of stress granules (Emara et al., 2010), suggesting that 5’ tRFs may be packaged in EVs as a result of cell stress, or cell activation, as in the case of T cells (Chiou et al., 2018). This process might be a step towards purging the excess cellular RNA or transferring specific messages to other cells. Interestingly, angiogenin, a ribonuclease activated by stress responses and responsible for the anticodon cleavage of tRNAs, has also been found in EVs, indicating that tRNA cleavage may also take place within EVs (Wei et al., 2017).

Various fragments of tRNA-Gly, tRNA-Asp, and tRNA-Glu were found within the most upregulated tRFs in both M1 and M2 EVs. Fragments of these tRNAs have previously been reported in EVs of various sources including cell lines (Sork et al., 2018, Wei et al., 2017), bone marrow- and adipose-mesenchymal stem cells (Baglio et al., 2015), placenta (Cooke et al., 2019), semen (Vojtech et al., 2014), plasma, serum, urine and bile (Srinivasan et al., 2019), and have been implicated in gene silencing by sequestering the Y-Box Binding Protein 1 (YBX1) (Goodarzi et al., 2015). Specifically, Goodarzi et al. (2015) showed that these YBX1-antagonistic tRFs dislocate YBX1 from the 3’untranslated region of oncogenic transcripts in breast cancer cells, leading to suppression of tumour growth and metastasis. Some of these tRF targets include the Eukaryotic translation initiation factor 4 gamma 1 (EIF4G1), the Integrin subunit beta 4 (ITGB4), and the Serine/threonine-protein kinase 1 (AKT1), which are involved in multiple cell functions outside of the context of cancer. Whether these tRFs can be delivered by EVs and exert their functions in recipient cells is unknown.

We report 34 tRFs that were significantly more abundant in M1 macrophage EVs compared to M2 EVs. These included 5’ end or internal fragments of tRNA-Leu(CAG, CAA, TAA), tRNA-Asp(GTC), tRNA-His(GTG), tRNA-Val(CAC and AAC), tRNA-Trp(CCA), tRNA-Met(CAT), tRNA-Gln(CTG), tRNA-Glu(TTC and CTC), tRNA-Gly(CCC), tRNA-Lys(CTT). Activated CD4^+^ T cells release EVs that are enriched in 5’tRFs derived from the tRNA-Leu (TAA and TAG), suggesting specific EV cargo loading in response to stimulation (Chiou et al., 2018), which might also be the case for macrophages. Li et al. (2016), showed that cleavage of tRNA-Val(CAC) by angiogenin was augmented during ischemic injury and hypoxia, and 5’ tRFs of this tRNA inhibit proliferation, migration and the tube formation capacity of endothelial cells, suggesting that the 5’tRF-Val(CAC) might have anti-angiogenic properties, in line with the characteristics of M1 macrophages. Fragments of the 5’end of tRNA-Gly(CCC),tRNA-Lys(CTT), and tRNA-Glu(CTC) favour respiratory syncytial virus (RSV) replication in cell lines by suppressing host defence genes like the apolipoprotein E receptor 2 (APOER2) (Wang et al., 2021, Deng et al., 2015). The functional implications of the overrepresentation of these tRFs in M1 macrophage EVs is yet to be investigated.

We found 32 tRFs to be significantly more abundant in M2 macrophage EVs compared to M1 EVs. These included 5’ end or internal fragments of tRNA-Glu(TTC and CTC), tRNA-Asp(GTC), tRNA-Arg(TCT, CCG, ACG), tRNA-Gly(GCC and CCC), tRNA-Lys(TTT), tRNA-Cys(GCA), tRNA-Phe(GAA), tRNA-Ile(AAT). Eight different itRFs arising from the tRNA-Glu(TTC) were found in M2 EVs, three of which (sequences ~ 36-69) were approximately 32-fold more abundant in M2 EVs. Different fragments of the tRNA-Glu(TTC) had previously been characterised as tumour-suppressors in gastric and thyroid cancers, likely involved in MAPK signalling (Xu et al., 2021, Shan et al., 2021).

Fragments of several lengths can be produced from the same tRNA. For example, in our dataset we observed various itRFs from the tRNA-His(GTG) including the sequences 2-24, 10-33, 15-34, etc. Each of these itRFs might be involved in different biological processes. This highlights the need for standardised nomenclature for the different tRFs to enable inter-study comparisons and metanalyses. The clinical significance of tRNAs and tRFs in EVs has recently been reviewed by Liu et al. (2022) and Weng et al. (2022).

In addition to miRNAs and tRFs, we also identified other small non-coding RNAs in M1 and M2 macrophage EVs, including Y-RNAs, followed by piRNAs, snRNAs and snoRNAs. Y-RNA derived small RNAs have been reported in EVs from various sources, including immune cells (Nolte-’t Hoen et al., 2012, Driedonks et al., 2018), cancer cells (Tosar et al., 2015, Lunavat et al., 2015), and body fluids (Yeri et al., 2017, Dhahbi et al., 2014, Vojtech et al., 2014), and sustain immunomodulatory properties (Haderk et al., 2017). More specifically, Y-RNA gene 4 (RNY4), present in the circulating EVs of patients with chronic lymphocytic leukemia, triggers the expression of the Programmed Death Ligand 1 (PDL-1) in monocytes, leading to immune-suppression (Haderk et al., 2017). In our dataset, RNY4 was more abundant in M2 macrophage EVs compared to M1 EVs, further supporting its anti-inflammatory role. PiR-33151 and piR-44757 were the two most abundant piRNAs in both M1 and M2 EVs. Upregulation of piR-33151 occurs in plasma and plasma EVs in patients with amyotrophic lateral sclerosis and chronic thromboembolic pulmonary hypertension, respectively (Joilin et al., 2020, Lipps et al., 2019). Little is known about the function of piR-44757, except for its possible role as an enhancer of the Myeloid differentiation primary response 88 (MYD88) expression, a protein adapter that facilitates signal transduction between immune cells (Fishilevich et al., 2017, Deguine and Barton, 2014). The most abundant snRNA in both M1 and M2 EVs was RNU2-1. Fragments of RNU2-1 are elevated in the blood of patients with central nervous system lymphomas (Baraniskin et al., 2016), melanoma (Kuhlmann et al., 2015), ovarian cancer (Kuhlmann et al., 2014), pancreatic and colorectal adenocarcinoma (Baraniskin et al., 2013), however the function of these RNAs and RNA fragments as well as their targets remain to be explored. Lastly, we identified several snoRNAs, including SNORDs, which drive methylation of rRNAs, SNORAs, which drive pseudo uridylation of rRNAs, and SCARNAs which facilitate methylation or pseudo uridylation of snRNAs and are localised to the Cajal body (Lafontaine and Tollervey, 1998, Darzacq et al., 2002). The snoRNA content of immune cell derived EVs changes in response to stimulatory or suppressive signals (Rimer et al., 2018). Specifically, activation of macrophages with lipopolysaccharide (LPS) results in accumulation of the SNORDs 32a, U33, U34, and U35a in secreted EVs, without accumulation in the cytoplasm, and uptake of these SNORDs occurs in distant tissues *in vivo,* resulting in modification/ methylation of their RNA targets (Rimer et al., 2018). We did not observe any statistically significant differences regarding these SNORDs between M1 and M2 EVs, which could reflect the different experimental settings; while we used a 10-day long polarisation protocol followed by a 24h EV generation period for human mo-macrophages, Rimer et al. (2018) collected EVs 1h after stimulation of mouse bone marrow derived macrophages and noticed clearance of these RNAs from the medium within 4h.

One limitation of small RNA sequencing (<75 nucleotides), and therefore our study, is the effect of the cut-off on detectable RNAs. For instance, snoRNAs with sizes ranging between 60 and 170 nucleotides might not be comprehensively represented in the final dataset. Thus, careful consideration is required when comparing EV RNA cargo information obtained using different sequencing methodologies. Analysing bulk EV RNA precludes the appreciation of the heterogeneity in the transcriptome of single EVs; detection of individual RNAs on a single EV basis, e.g., by flow cytometry, could be employed to study immune cell EV RNA cargo at a more granular level. To date, both small and total RNA sequencing have considerably advanced our understanding of the diverse EV RNA cargo (reviewed by Dellar et al. (2022)).

In this study, we isolated EVs from M1 -pro-inflammatory- and M2 -anti-inflammatory-primary macrophages using size exclusion chromatography and performed small RNA sequencing to provide a comprehensive analysis of their small non-coding RNA cargo. Overall, we propose that the M1/M2 EV transcriptome is shaped by distinct non-coding RNA structures. Within each subtype of small RNA, we demonstrate many significant differences between M1 and M2 EVs, likely aided by our long polarisation protocol and the absence of any serum at any stage of culture, which could affect cell function and contaminate EV isolates. Functional enrichment analysis of miRNA targets revealed contrasting targets of EVs from M1 and M2 macrophages, which likely contribute to their functional effects on their environment. Similarly, differentially sorted tRNA fragments, piRNA, snRNA, snoRNA and Y-RNAs in M1 and M2 EVs might exert functional effects in recipient cells, or their secretion might be important for the parental cell. A deeper look into the relevance of small RNA fragments transported via EVs is an important next step.

## Data availability statement

All the sequencing data generated in this study have been deposited to the Gene Expression Omnibus (GEO) repository database with the accession number GSE207286. Also, this study is registered on EV-TRACK with reference number EV220120.

## Acknowledgements

Research reported in this publication was supported by National Institute for Child Health (NICHD) of the National Institutes of Health under award number R01HD093801. The content is solely the responsibility of the authors and does not necessarily represent the official views of the National Institutes of Health. MV is supported by the EU grant No. 964712 (EU-H2020-FETOpen PRIME).

## Author contributions

Paschalia Pantazi: experimental work, data curation, writing – original draft. Toby Clements: experimental work, writing – review & editing. Morten Venø: data analysis, writing – review & editing. Vikki Abrahams: methodology, writing – review & editing. Beth Holder: funding acquisition, supervision, methodology, data curation, writing – original draft and review & editing. All authors have read and approved the final version of the manuscript.

