## Supplemental Table 2 for "Distinct non-coding RNA cargo of extracellular vesicles from M1 and M2 human primary macrophages"

**Supplemental Table 2.** Significantly differentially abundant miRNAs measured in M1 versus M2 macrophage EVs. Log2FC; Log2 fold change.

| Higher in M1 macrophage EVs | | | |
| --- | --- | --- | --- |
| microRNA name | **miRBase accession** | **log2FC** | **adjusted p value** |
| hsa-miR-125a-5p | MIMAT0000443 | -2.19 | 8.44E-17 |
| hsa-miR-23b-3p | MIMAT0000418 | -2.44 | 2.03E-12 |
| hsa-miR-146b-5p | MIMAT0002809 | -2.93 | 1.03E-10 |
| hsa-let-7i-5p | MIMAT0000415 | -2.23 | 7.00E-10 |
| hsa-miR-146b-3p | MIMAT0004766 | -2.53 | 3.77E-09 |
| hsa-miR-99b-5p | MIMAT0000689 | -1.49 | 4.85E-09 |
| hsa-miR-1307-3p | MIMAT0005951 | -1.58 | 2.82E-08 |
| hsa-miR-147b-3p | MIMAT0004928 | -4.12 | 1.26E-07 |
| hsa-miR-222-3p | MIMAT0000279 | -2.13 | 3.24E-07 |
| hsa-miR-99b-3p | MIMAT0004678 | -2.87 | 5.05E-06 |
| hsa-let-7e-5p | MIMAT0000066 | -2.08 | 1.53E-05 |
| hsa-miR-210-3p | MIMAT0000267 | -2.91 | 2.75E-05 |
| hsa-miR-9-5p | MIMAT0000441 | -2.06 | 8.66E-05 |
| hsa-miR-511-5p | MIMAT0002808 | -3.36 | 1.60E-04 |
| hsa-miR-146a-5p | MIMAT0000449 | -1.76 | 2.57E-04 |
| hsa-miR-155-5p | MIMAT0000646 | -2.00 | 2.73E-04 |
| hsa-miR-187-3p | MIMAT0000262 | -4.54 | 5.50E-04 |
| hsa-miR-671-5p | MIMAT0003880 | -2.16 | 5.97E-04 |
| hsa-miR-378c | MIMAT0016847 | -1.59 | 6.73E-04 |
| hsa-miR-7-5p | MIMAT0000252 | -1.60 | 8.69E-04 |
| hsa-miR-221-5p | MIMAT0004568 | -2.72 | 1.18E-03 |
| hsa-miR-193a-5p | MIMAT0004614 | -1.92 | 2.58E-03 |
| hsa-miR-22-3p | MIMAT0000077 | -1.71 | 2.58E-03 |
| hsa-miR-9985 | MIMAT0039763 | -2.85 | 3.55E-03 |
| hsa-miR-218-5p | MIMAT0000275 | -2.97 | 4.54E-03 |
| hsa-miR-132-5p | MIMAT0004594 | -2.31 | 5.24E-03 |
| hsa-miR-92b-5p | MIMAT0004792 | -3.24 | 5.37E-03 |
| hsa-miR-181b-5p | MIMAT0000257 | -1.23 | 8.42E-03 |
| hsa-miR-24-3p | MIMAT0000080 | -1.04 | 1.02E-02 |
| hsa-miR-125a-3p | MIMAT0004602 | -2.74 | 1.25E-02 |
| hsa-miR-4773 | MIMAT0019928 | -3.39 | 1.61E-02 |
| hsa-miR-3614-5p | MIMAT0017992 | -2.81 | 1.86E-02 |
| hsa-let-7i-3p | MIMAT0004585 | -2.06 | 2.48E-02 |
| hsa-miR-181d-5p | MIMAT0002821 | -2.07 | 2.48E-02 |
| hsa-miR-4677-3p | MIMAT0019761 | -2.73 | 2.91E-02 |
| hsa-miR-1843 | MIMAT0039764 | -1.72 | 3.07E-02 |
| hsa-miR-181a-5p | MIMAT0000256 | -0.82 | 3.25E-02 |
| hsa-miR-22-5p | MIMAT0004495 | -1.35 | 3.31E-02 |
| hsa-miR-212-5p | MIMAT0022695 | -2.12 | 3.42E-02 |
| hsa-miR-505-3p | MIMAT0002876 | -1.40 | 3.75E-02 |
| hsa-miR-491-5p | MIMAT0002807 | -1.68 | 3.89E-02 |
| hsa-miR-107 | MIMAT0000104 | -0.93 | 4.03E-02 |
| hsa-let-7d-5p | MIMAT0000065 | -0.84 | 4.40E-02 |
| Higher in M2 macrophage EVs | | | |
| microRNA name | **miRBase accession** | **log2FC** | **adjusted p value** |
| hsa-miR-205-5p | MIMAT0000266 | 3.49 | 1.54E-07 |
| hsa-miR-96-5p | MIMAT0000095 | 3.70 | 2.56E-04 |
| hsa-miR-625-3p | MIMAT0004808 | 3.20 | 9.66E-04 |
| hsa-miR-432-5p | MIMAT0002814 | 4.87 | 3.55E-03 |
| hsa-miR-20a-5p | MIMAT0000075 | 1.16 | 4.23E-03 |
| hsa-miR-126-5p | MIMAT0000444 | 3.00 | 5.24E-03 |
| hsa-miR-182-5p | MIMAT0000259 | 2.74 | 6.64E-03 |
| hsa-miR-93-5p | MIMAT0000093 | 1.36 | 8.42E-03 |
| hsa-miR-345-5p | MIMAT0000772 | 1.29 | 1.11E-02 |
| hsa-miR-183-5p | MIMAT0000261 | 2.89 | 1.26E-02 |
| hsa-miR-1287-5p | MIMAT0005878 | 2.01 | 1.39E-02 |
| hsa-miR-4433b-5p | MIMAT0030413 | 4.34 | 1.60E-02 |
| hsa-miR-382-5p | MIMAT0000737 | 3.89 | 1.86E-02 |
| hsa-miR-95-3p | MIMAT0000094 | 1.65 | 1.86E-02 |
| hsa-miR-92a-3p | MIMAT0000092 | 1.61 | 2.11E-02 |
| hsa-miR-126-3p | MIMAT0000445 | 2.69 | 2.47E-02 |
| hsa-miR-1249-3p | MIMAT0005901 | 2.18 | 2.47E-02 |
| hsa-let-7d-3p | MIMAT0004484 | 1.45 | 2.48E-02 |
| hsa-miR-145-5p | MIMAT0000437 | 4.16 | 2.48E-02 |
| hsa-miR-451a | MIMAT0001631 | 3.63 | 2.48E-02 |
| hsa-miR-486-5p | MIMAT0002177 | 4.04 | 2.48E-02 |
| hsa-miR-16-5p | MIMAT0000069 | 0.91 | 2.54E-02 |
| hsa-miR-1268a | MIMAT0005922 | 2.18 | 2.91E-02 |
| hsa-miR-1294 | MIMAT0005884 | 3.74 | 2.91E-02 |
| hsa-miR-151a-3p | MIMAT0000757 | 1.58 | 2.91E-02 |
| hsa-miR-431-5p | MIMAT0001625 | 4.10 | 2.97E-02 |
| hsa-miR-223-3p | MIMAT0000280 | 1.16 | 3.00E-02 |
| hsa-miR-199a-3p | MIMAT0000232 | 1.47 | 3.01E-02 |
| hsa-miR-193b-5p | MIMAT0004767 | 2.25 | 4.03E-02 |
