## Supplemental Table 3 for "Distinct non-coding RNA cargo of extracellular vesicles from M1 and M2 human primary macrophages"

**Supplemental Table 3.** The top 20 targets of M1 and M2 EVs determined using MEINTURNET analysis, and the miRNAs targeting these.

| Gene Symbol | p-value | FDR | microRNA 1 | microRNA 2 | microRNA 3 | microRNA 4 | microRNA 5 | microRNA 6 | microRNA 7 | microRNA 8 | microRNA 9 | microRNA 10 |
| --- | --- | --- | --- | --- | --- | --- | --- | --- | --- | --- | --- | --- |
| SORT1 | 1.59E-07 | 6.41E-04 | hsa-miR-222-3p | hsa-miR-218-5p | hsa-miR-181d-5p | hsa-miR-181b-5p | hsa-miR-181a-5p | - | - | - | - | - |
| LBR | 4.22E-07 | 6.41E-04 | hsa-miR-181a-5p | hsa-miR-24-3p | hsa-let-7d-5p | hsa-miR-23b-3p | hsa-miR-107 | hsa-miR-146b-5p | hsa-miR-146a-5p | hsa-miR-181d-5p | hsa-miR-181b-5p | - |
| TFRC | 4.22E-07 | 6.41E-04 | hsa-miR-22-3p | hsa-miR-210-3p | hsa-miR-125a-5p | hsa-miR-9-5p | hsa-miR-218-5p | hsa-miR-7-5p | hsa-miR-181d-5p | hsa-miR-181b-5p | hsa-miR-181a-5p | - |
| CHD9 | 4.31E-07 | 6.41E-04 | hsa-miR-210-3p | hsa-miR-155-5p | hsa-miR-7-5p | hsa-let-7e-5p | hsa-miR-22-3p | hsa-miR-181a-5p | hsa-miR-181b-5p | hsa-miR-181d-5p | - | - |
| IL6 | 5.62E-07 | 6.68E-04 | hsa-miR-155-5p | hsa-miR-107 | hsa-miR-146b-5p | hsa-miR-9-5p | hsa-miR-146a-5p | hsa-miR-125a-3p | - | - | - | - |
| GANAB | 8.69E-07 | 8.62E-04 | hsa-miR-23b-3p | hsa-miR-222-3p | hsa-miR-181b-5p | hsa-miR-181a-5p | hsa-miR-107 | hsa-miR-155-5p | hsa-miR-146b-3p | - | - | - |
| EGFR | 1.52E-06 | 1.29E-03 | hsa-miR-7-5p | hsa-miR-146a-5p | hsa-miR-155-5p | hsa-miR-125a-5p | hsa-miR-218-5p | hsa-miR-491-5p | hsa-miR-146b-5p | - | - | - |
| EDEM3 | 1.78E-06 | 1.32E-03 | hsa-miR-155-5p | hsa-miR-218-5p | hsa-miR-505-3p | hsa-let-7e-5p | hsa-let-7d-5p | hsa-let-7i-5p | hsa-miR-146b-5p | hsa-miR-146a-5p | hsa-miR-9-5p | - |
| MTUS1 | 2.54E-06 | 1.53E-03 | hsa-miR-125a-5p | hsa-let-7i-5p | hsa-let-7d-5p | hsa-let-7e-5p | hsa-miR-181d-5p | hsa-miR-181a-5p | hsa-miR-181b-5p | - | - | - |
| FUBP1 | 2.58E-06 | 1.53E-03 | hsa-miR-155-5p | hsa-miR-222-3p | hsa-miR-22-3p | hsa-miR-218-5p | hsa-let-7i-3p | - | - | - | - | - |
| MEG3 | 3.23E-06 | 1.74E-03 | hsa-miR-181a-5p | hsa-miR-181b-5p | hsa-miR-181d-5p | hsa-miR-125a-5p | - | - | - | - | - | - |
| SLC10A7 | 4.54E-06 | 2.25E-03 | hsa-let-7d-5p | hsa-miR-218-5p | hsa-let-7e-5p | hsa-let-7i-5p | hsa-miR-222-3p | hsa-miR-181d-5p | hsa-miR-181a-5p | hsa-miR-181b-5p | - | - |
| RP2 | 5.74E-06 | 2.63E-03 | hsa-miR-155-5p | hsa-miR-181d-5p | hsa-miR-181b-5p | hsa-miR-181a-5p | - | - | - | - | - | - |
| MYC | 9.21E-06 | 3.03E-03 | hsa-miR-24-3p | hsa-miR-155-5p | hsa-miR-7-5p | hsa-miR-4677-3p | hsa-miR-222-3p | hsa-miR-125a-3p | hsa-miR-125a-5p | hsa-let-7i-5p | hsa-let-7e-5p | hsa-let-7d-5p |
| SSSCA1 | 9.46E-06 | 3.03E-03 | hsa-miR-222-3p | hsa-miR-155-5p | hsa-miR-24-3p | hsa-miR-221-5p | - | - | - | - | - | - |
| NR6A1 | 9.83E-06 | 3.03E-03 | hsa-miR-9-5p | hsa-miR-181a-5p | hsa-miR-218-5p | hsa-let-7e-5p | hsa-let-7i-5p | hsa-let-7d-5p | hsa-miR-181d-5p | hsa-miR-181b-5p | - | - |
| AKAP8 | 1.05E-05 | 3.03E-03 | hsa-miR-218-5p | hsa-let-7d-5p | hsa-let-7i-5p | hsa-let-7e-5p | hsa-miR-146b-5p | hsa-miR-146a-5p | - | - | - | - |
| MTOR | 1.14E-05 | 3.03E-03 | hsa-miR-99b-5p | hsa-miR-181b-5p | hsa-miR-218-5p | hsa-miR-193a-5p | hsa-miR-125a-5p | - | - | - | - | - |
| CXCL8 | 1.25E-05 | 3.03E-03 | hsa-miR-146a-5p | hsa-miR-155-5p | hsa-let-7d-5p | hsa-let-7e-5p | hsa-let-7i-5p | hsa-miR-4677-3p | - | - | - | - |
| FAT1 | 1.34E-05 | 3.03E-03 | hsa-miR-222-3p | hsa-miR-181a-5p | hsa-miR-218-5p | - | - | - | - | - | - | - |
