## Supplemental Table 5 for "Distinct non-coding RNA cargo of extracellular vesicles from M1 and M2 human primary macrophages"

**Supplemental Table 2.** Significantly differentially abundant tRNA fragments measured in M1 versus M2 macrophage EVs. Log2FC; Log2 fold change.

| Higher in M1 macrophage EVs | | |
| --- | --- | --- |
| MINTbase ID | **log2FC** | **adjusted p value** |
| tRF-20-WR381H93 | -3.36 | 1.14E-05 |
| tRF-39-S5QKF1R3WE8RO8IS | -3.54 | 2.55E-05 |
| tRF-17-RXSINHQ | -2.35 | 2.55E-05 |
| tRF-19-O7M8LO1Z | -1.82 | 7.28E-05 |
| tRF-32-897PVP941QKSJ | -2.08 | 9.84E-05 |
| tRF-20-WRD81H93 | -3.78 | 8.32E-04 |
| tRF-20-WRD81H93 | -3.78 | 8.32E-04 |
| tRF-22-MUWLV47PO | -4.51 | 1.17E-03 |
| tRF-34-79MP9PMNH5IS15 | -1.87 | 2.95E-03 |
| tRF-31-821IQ4YUH5S80 | -4.55 | 3.43E-03 |
| tRF-17-F8DHXY4 | -2.22 | 3.70E-03 |
| tRF-23-VP9N15WV0E | -3.12 | 4.31E-03 |
| tRF-28-PSQP4PW3FJD0 | -1.66 | 6.31E-03 |
| tRF-37-Q2FWQ8P6U1J0SUP | -4.27 | 7.09E-03 |
| tRF-23-SERXPIN2D5 | -3.20 | 8.86E-03 |
| tRF-19-SP58301Z | -1.40 | 8.98E-03 |
| tRF-20-SERXPIN2 | -4.49 | 9.91E-03 |
| tRF-19-9N15WV2P | -4.41 | 9.91E-03 |
| tRF-18-0L9RQ4DZ | -2.11 | 9.91E-03 |
| tRF-34-Q99P9P9NH57S15 | -2.28 | 1.06E-02 |
| tRF-19-OQIMLWIZ | -2.97 | 1.60E-02 |
| tRF-19-821IQ4KK | -2.73 | 1.60E-02 |
| tRF-16-YU4RRW0 | -1.86 | 1.60E-02 |
| tRF-31-897PVP941QKSD | -1.82 | 1.60E-02 |
| tRF-36-897PVP941QKS3WD | -1.91 | 1.69E-02 |
| tRF-20-897PVP94 | -2.47 | 2.28E-02 |
| tRF-26-L725QB1MK8E | -3.52 | 2.56E-02 |
| tRF-19-73VL4YHE | -2.62 | 2.67E-02 |
| tRF-37-YRI7XUK87SIRMJ4 | -2.39 | 2.67E-02 |
| tRF-32-79MP9P9MH57SJ | -1.48 | 2.90E-02 |
| tRF-33-PS5U8918W8RVY | -1.50 | 3.33E-02 |
| tRF-18-9N15WV0E | -1.95 | 3.37E-02 |
| tRF-38-S5QKF1R3WE8RO8DX | -3.97 | 4.60E-02 |
| tRF-29-389MV47P59I8 | -2.06 | 4.77E-02 |
| Higher in M2 macrophage EVs | | |
| MINTbase ID | **log2FC** | **adjusted p value** |
| tRF-33-0668K87SERM404 | 5.58 | 8.65E-09 |
| tRF-32-KPQJ3KYU4RRW4 | 5.40 | 4.73E-06 |
| tRF-33-EH623K7SIR3DDY | 1.79 | 6.32E-06 |
| tRF-30-JJ6RRNLIK898 | 4.48 | 9.80E-06 |
| tRF-41-L7S5QKF1R3WE8RO80 | 3.90 | 1.23E-05 |
| tRF-19-R118LOJX | 3.16 | 1.25E-05 |
| tRF-33-EH623K76IR3DDY | 1.59 | 2.70E-03 |
| tRF-39-3W2VR008R959KU1Y | 3.27 | 3.31E-03 |
| tRF-16-7SBRM50 | 2.35 | 3.43E-03 |
| tRF-17-PV6RRNK | 3.18 | 5.06E-03 |
| tRF-29-1HPSR9O93324 | 3.04 | 6.04E-03 |
| tRF-17-7SBRM54 | 1.63 | 6.20E-03 |
| tRF-37-87R8WP9N1EWJQ72 | 2.21 | 7.65E-03 |
| tRF-38-87R8WP9N1EWJQ7D1 | 2.49 | 9.91E-03 |
| tRF-16-7S1RMH0 | 1.54 | 9.91E-03 |
| tRF-32-31QJ3KYUYRR64 | 2.18 | 1.09E-02 |
| tRF-35-86V8WPMN1E8Y7Z | 1.67 | 1.09E-02 |
| tRF-30-87R8WP9N1EWJ | 1.32 | 1.14E-02 |
| tRF-19-R95933FY | 2.46 | 2.17E-02 |
| tRF-16-PYRP4PE | 1.43 | 2.17E-02 |
| tRF-19-OR183OJX | 2.72 | 2.20E-02 |
| tRF-17-R95M3J4 | 1.41 | 2.28E-02 |
| tRF-19-69M8LOJX | 1.73 | 2.53E-02 |
| tRF-31-QKF1R3WE8RO80 | 1.26 | 2.67E-02 |
| tRF-35-87R8WP9I1EWJQZ | 2.49 | 3.18E-02 |
| tRF-32-L85DMKYUYRLH4 | 1.67 | 3.33E-02 |
| tRF-16-76DRR4D | 1.57 | 3.45E-02 |
| tRF-16-76DRR4D | 1.57 | 3.45E-02 |
| tRF-17-SR9NLMJ | 3.01 | 3.71E-02 |
| tRF-36-Q1Q89P9L8422YRB | 2.36 | 3.93E-02 |
| tRF-39-87R8WP9N1EWJQ7FV | 1.94 | 4.55E-02 |
| tRF-34-PNR8YP9LON4VHM | 1.11 | 4.88E-02 |
